# Uncovering Cas9 PAM diversity through metagenomic mining and machine learning

**DOI:** 10.1101/2025.08.13.668647

**Authors:** Tao Fang, Lea Bogensperger, Lilith Feer, Ahmed Allam, Valentyn Bezshapkin, Zsolt Balázs, Christian von Mering, Shinichi Sunagawa, Michael Krauthammer, Gerald Schwank

**Author notes:** Corresponding authors Correspondence to Michael Krauthammer or Gerald Schwank. These authors contributed equally (Tao Fang, Lea Bogensperger).

## Abstract

Recognition of protospacer adjacent motifs (PAMs) is crucial for target site recognition by CRISPR–Cas systems. In genome editing applications, the requirement for specific PAM sequences at the target locus imposes substantial constraints, driving efforts to search for novel Cas9 orthologs with extended or alternative PAM compatibilities. Here, we present CRISPR-PAMdb, a comprehensive and publicly accessible database compiling Cas9 protein sequences from 3.8 million bacterial and archaeal genomes and PAM profiles from 7.4 million phage and plasmid sequences. Through spacer–protospacer alignment, we inferred consensus PAM preferences for 8,003 unique Cas9 clusters. To extend PAM discovery beyond traditional alignment-based approaches, we developed CICERO, a machine learning model predicting PAM preferences directly from Cas9 protein sequences. Built on the ESM2 protein language model and trained on the CRISPR–PAMdb database, CICERO achieved an average accuracy of 0.68 on test data and 0.75 on experimentally validated Cas9 orthologs. For Cas9 clusters where alignment-based predictions were infeasible, CICERO generated PAM profiles for an additional 50,308 Cas9 proteins, including 17,453 high-confidence predictions with accuracies above 0.86. CRISPR–PAMdb, alongside CICERO models, enables large-scale exploration of PAM diversity across Cas9 proteins, accelerating design of next-generation CRISPR-Cas9 tools for precise genome engineering.

## Introduction

CRISPR–Cas systems, which provide adaptive immunity in prokaryotes, have been repurposed as powerful tools for modern genome editing. In CRISPR-Cas systems, such as type II (Cas9) and type V (Cas12), the recognition of protospacer adjacent motifs (PAMs)—short DNA sequences (usually 2–6 base pairs) adjacent to the target site—is critical. The PAM acts as a molecular signal, enabling Cas proteins to bind specific DNA sequences and distinguish between self and non-self DNA, thus preventing unintended cleavage of the host genome. Upon PAM recognition, Cas unwinds adjacent double-stranded DNA, allowing R-loop formation where the CRISPR RNA (crRNA) pairs with the target strand. If the crRNA spacer is perfectly complementary to the target sequence, full base pairing occurs and the Cas protein cleaves the DNA, enabling immune defense or genome editing^1–3^.

While critical for precision, PAM recognition imposes a major constraint on CRISPR-Cas systems by limiting the range of targetable genomic sites. This restriction presents a significant challenge for genome editing, particularly in therapeutic contexts where flexible targeting is crucial for treating diverse genetic disorders^4^. Expanding PAM diversity, either through protein engineering or the discovery of natural Cas proteins with novel PAM specificities, is therefore a key priority for increasing the versatility and effectiveness of CRISPR-Cas technologies^5–7^.

To uncover new CRISPR-Cas systems with novel PAM specificities, recent studies have leveraged metagenomic data from environmental microbial communities. Ciciani et al. developed a computational pipeline that analyzes metagenome and virome assemblies and uses spacer–protospacer alignment to infer PAM preferences for Cas9 proteins^8^. Similarly, Ruffolo et al. also adopted spacer–protospacer alignment, but utilized more comprehensive metagenomic and phage datasets, allowing them to infer PAM specificities for a larger number of Cas9 proteins^9,10^.

However, a key limitation of both studies is that their databases are not publicly accessible and thus cannot be employed for the development of CRISPR–Cas9 systems with novel PAM recognition. Moreover, a major bottleneck in the metagenomic mining approach is the limited number of protospacer matches per Cas9 cluster in phage and plasmid databases, leaving the majority of Cas9 clusters without assignable PAM preferences. To address this gap, we introduce CRISPR-PAMdb, a publicly available, curated database of Cas9 protein sequences and their inferred PAMs, derived from mining over 3.8 million bacterial and archaeal genomes and more than 7.4 million phage and plasmid sequences. Inferred PAMs are associated with individual Cas9 proteins by aligning CRISPR spacers to protospacers within phage and plasmid sequences, followed by the extraction of consensus PAM sequences. Furthermore, we present CICERO (CRISPR Cas9 PAM predictor model), a machine learning model that is trained on the CRISPR-PAMdb database and capable of predicting PAM preferences directly from Cas9 protein sequences using an ESM2 protein language model backbone^11^. By providing these resources and the accompanying pipelines, we aim to facilitate broader exploration of the Cas9 family to accelerate rational design of next-generation genome-editing tools with alternative PAM preferences.

## Results

### Alignment-Based Pipeline for CRISPR-Cas9 System Mining and PAM Inference

We first developed a computational pipeline (Fig. 1A and Methods section) to systematically mine CRISPR-Cas systems from bacterial and archaeal (meta)genomes, along with their associated protospacers from phage and plasmid sequences. Due to computational constraints, we focused on identifying CRISPR-Cas9 systems, although the pipeline is also adaptable to other CRISPR-Cas systems. The pipeline first retrieved 3,747,151 bacterial and archaeal genomes from the mOTUs4 database, consisting of isolate genomes, single amplified genomes, and metagenome-assembled genomes^12^, and an additional 25,371 archaeal genomes from the European Nucleotide Archive (ENA)^13^. From these, we identified CRISPR arrays, including repeat and spacer sequences, and flanking protein-coding genes. These genes were subsequently screened using curated Cas hidden Markov models^14^, resulting in the identification of 21,265,037 Cas proteins and 430,685 Cas9 proteins. Based on the taxonomy information from mOTUs4, Cas proteins were identified in 1,658,140 bacterial genomes, of which 335,447 (20%) contain Cas9 proteins. In contrast, among the 23,352 archaeal genomes with Cas proteins, only 429 contain Cas9 proteins (Fig. 1B), consistent with previous observations of Cas9 rarity in archaea. This scarcity may reflect lower viral pressure in archaeal environments, favoring alternative defense mechanisms. Alternatively, it could result from a higher fitness cost of maintaining Cas9 in energy-limited or extreme environments^15–18^.

**Figure 1.**
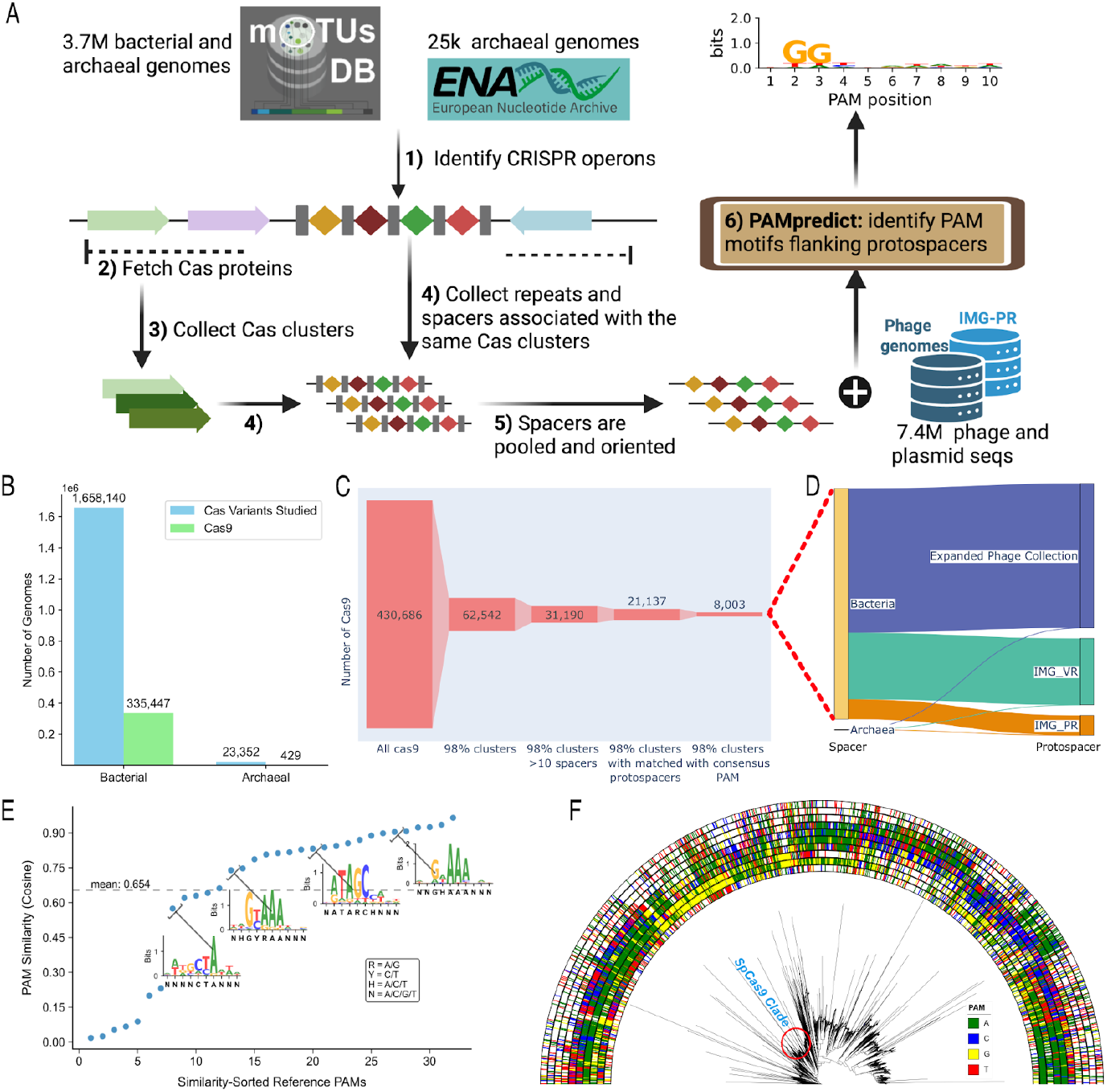
(A) Schematic overview of our mining pipeline. 1) Identify CRISPR repeat-spacer arrays from bacterial and archaeal genomes; 2) Fetch Cas proteins; 3) Collect Cas clusters; 4) Collect repeats and spacers associated with the same Cas clusters; 5) Spacers are pooled and oriented consistently by clustering repeats; 6) Map pooled spacers to identify PAMs flanking protospacers in dereplicated phage and plasmid sequences using PAMpredict^8^. (B) Count and sources of mined Cas proteins. (C) Count of Cas9 variants at each pipeline stage. (D) The Sankey plot illustrates the mapping between spacers in the bacterial and archaeal genomes and their mapped protospacers in phage and plasmid sequences for the final set of 8,003 Cas9 variants with consensus PAM inferences. (E) Similarity between 32 reference PAM sequences with consensus PAM inferences derived from their most similar Cas9 protein clusters (≥98% sequence similarity). Accompanying sequence logo plots illustrate the inferred PAM profiles across different levels of PAM similarity. The string shown below the x-axis in each plot represents the corresponding reference PAM sequence. (F) Phylogenetic tree of Cas9 protein clusters sharing 98% sequence similarity. Sequences were aligned using MAFFT^22^, the tree was built with FastTree^23^, and visualized in iTOL^24^. Annotations, from inner to outer rings, indicate the most likely nucleotide inferred at each of the 10 PAM positions.

To identify the protospacer adjacent motifs (PAMs) associated with the Cas9 family, we grouped crRNAs (spacers) from 62,542 Cas9 protein clusters sharing ≥98% sequence identity. Among these, 31,190 clusters contained more than 10 spacers. For 21,137 of them, we successfully mapped protospacers within 7,413,108 phage and plasmid sequences derived from a comprehensive collection of mobile genetic elements, including plasmid sequences from IMG-PR^19^, phage sequences from IMG-VR^20^, and additional phage sequences from an expanded collection compiled from other studies (see Methods section for details). Importantly, the alignment of spacers to their corresponding protospacers allowed us to infer PAM preferences for 8,003 Cas9 clusters using PAMpredict^8^ (Fig. 1C). Of note, the accuracy of spacer–protospacer-based PAM inference strongly depends on the availability of unique protospacers in phage sequences. We therefore evaluated the contribution of our collected mobile genetic elements database compared to the commonly used viral genome database, IMG-VR^20^. Interestingly, we found that over half of the mapped protospacers were unique to the mobile genetic elements dataset (Fig. 1D). This highlights the importance of expanding reference datasets for PAM discovery through spacer-protospacer alignment and suggests limitations in the field’s reliance on IMG-VR and IMG-PR as standard references.

Next, we evaluated the performance of our pipeline and benchmarked the inferred PAMs using 79 Cas9 proteins for which the PAM profiles were experimentally characterized^21^. Our mined Cas9 dataset included 66 of the 79 reference Cas9 proteins when using a 98% threshold for sequence similarity. After filtering, consensus PAM inferences were obtained for 32 out of the 66 Cas9 clusters (Supplementary Fig. 1B). We then compared the experimentally determined PAM profiles of the reference Cas9 variants with the bioinformatically inferred PAM profiles using augmented cosine similarity. Confirming the high accuracy of our bioinformatic PAM identification approach, the majority of the 32 Cas9 clusters showed cosine similarities above 0.5 to the reported PAM profile (average cosine similarity = 0.654; Fig. 1E). Next, we constructed a phylogenetic tree of all 8,003 identified Cas9 protein clusters with their consensus PAMs. This tree reveals that closely related Cas9 sequences typically detect similar PAMs. Moreover, it highlights the large diversity of PAMs within the Cas9 family, although purines are favored in the PAM sequences of most orthologs (Fig. 1F).

### Machine Learning-Based PAM Prediction for Cas9 Proteins using CICERO

For the majority of the 62,542 identified Cas9 protein clusters, we were unable to map a sufficient number of protospacers in phage and plasmid databases to infer PAMs. To overcome this limitation, we next developed and trained CICERO (<u>C</u>R<u>I</u>SPR <u>C</u>as9 PAM predictor model), a machine learning model that is capable of predicting PAM profiles directly from CRISPR Cas9 sequences using a protein language model (pLM) backbone (ESM2)^11^. Our training pipeline, similar to Nayfach et al.^9^, follows a two-phase scheme (Fig. 2A). In phase 1, CICERO is trained to predict PAMs directly for given Cas9 proteins as inputs. Thereby, the protein language model embeddings are extended by a multi-layer perceptron (MLP) network to predict PAM profiles, which reflect how likely each nucleotide is to appear at each position (see *Methods* for more technical details). In phase 2, which only takes place after phase 1 is completed, the setup is further refined to have another MLP layer on top of the trained model from phase 1 to predict corresponding confidence estimates of the predictions. Essentially, this confidence prediction model learns to estimate the accuracy of PAM predictions from phase 1, thereby providing insight into the reliability of each prediction.

**Figure 2.**
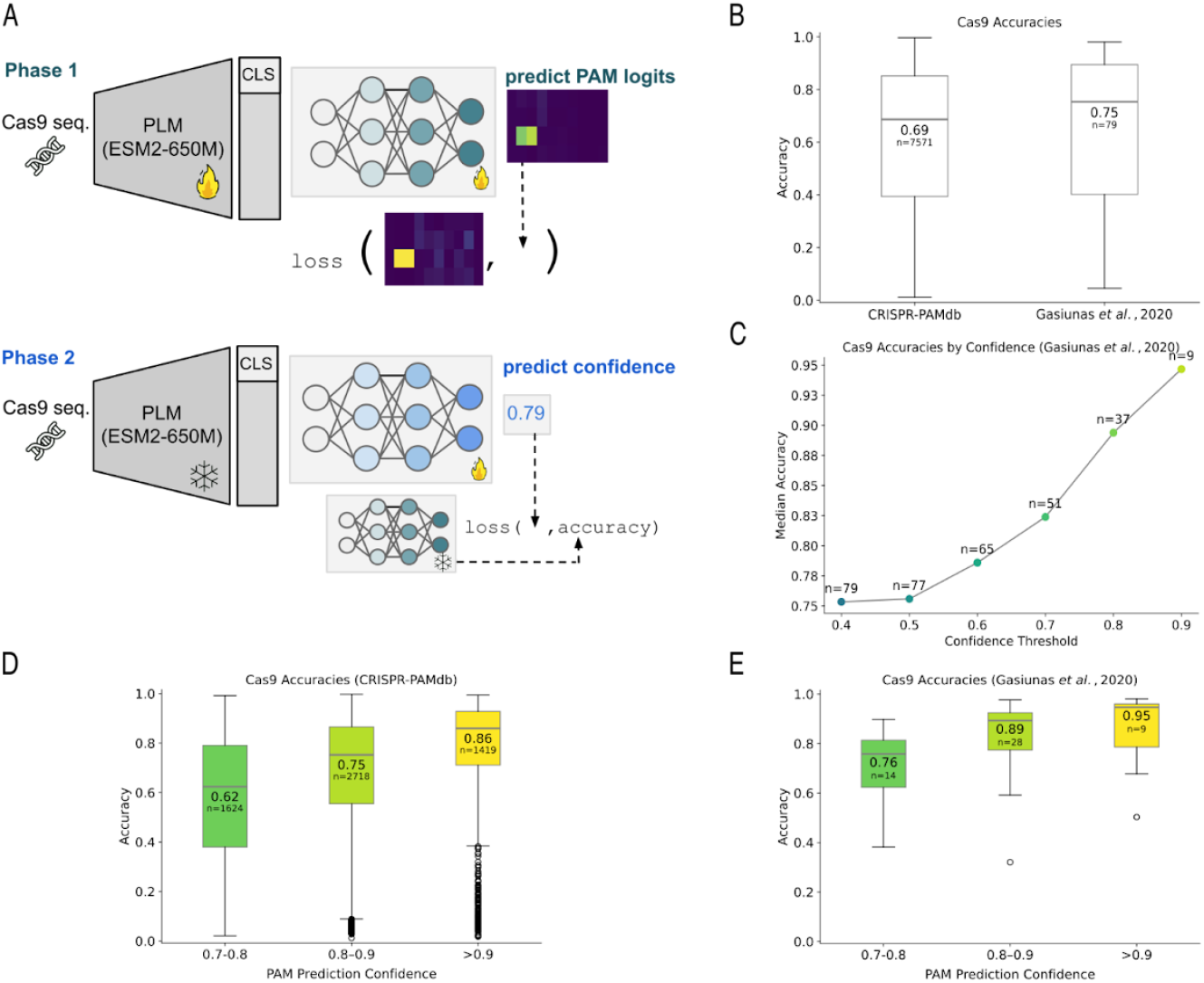
(A) Two-phase training pipeline similar to Nayfach et al.^9^ using a protein language model as backbone. In phase 1, the model is trained to predict PAM profiles given Cas9 input sequences. Phase 2 extends upon this by subsequent training of a confidence prediction module, which is trained to regress the PAM prediction accuracy, computed as augmented cosine similarity to the target PAM profile. (B) Median accuracies on CRISPR-PAMdb Cas9 sequences aggregated over all cross-validation folds as well as on an external validation set (Gasiunas et al.^21^). Accuracy is computed as augmented cosine similarity between predicted and inferred PAMs using alignment-based methods or experimentally validated PAMS from Gasiunas et al.^21^. (C) Median accuracy for the 79 external validation set, filtered according to varying confidence thresholds. Retaining only predictions with a confidence of at least 0.8 yields a much higher accuracy of 0.89 (for n=37 filtered samples) compared to the baseline accuracy of 0.75 based on all 79 sequences. (D) Median accuracies on CRISPR-PAMdb Cas9 sequences aggregated over all cross-validation folds as in (B) with additional grouping by predicted confidence. The number of samples in each confidence bin is shown, out of a total of 7,571 Cas9 samples. (E) Median accuracies on the external validation set (Gasiunas et al.^21^) as in (B) with additional grouping by predicted confidence. The number of samples in each confidence bin is shown, out of a total of 79 Cas9 samples.

We trained the final model, CICERO-650M, using five-fold cross-validation on the CRISPR-PAMdb dataset with the 8,003 Cas9 clusters with inferred PAM preference (see Methods section for details). When comparing predicted to inferred PAM profiles using augmented cosine similarity, CICERO-650M achieved a median accuracy of 0.69 ± 0.03 across the test splits (i.e., unseen data) (Fig. 2B; Supplementary Fig. 2 for qualitative results). Furthermore, performance on an independent benchmark of 79 experimentally validated Cas9 proteins yielded a median accuracy of 0.75 (Fig. 2B; Supplementary Fig. 3).

We further tested the utility of the trained confidence network (i.e., confidence head) on top of the PAM prediction system by generating confidence scores. The confidence predictions on the 79 external validation data are shown to strongly correlate with the true prediction accuracy with r = 0.8 (p = 8.8e-19), enabling a simple thresholding strategy where only predictions above a certain confidence threshold are accepted. This strategy substantially increased the median prediction accuracy for predictions with higher confidence thresholds (Fig. 2C). Thus, restricting outputs to high-confidence predictions leads to improved reliability. Grouping PAM predictions for the 8,003 Cas9 clusters with inferred PAM preference (over all five folds of CRISPR-PAMdb test data) into confidence bins (0.7-0.8, 0.8-0.9,>0.9) therefore resulted in substantially higher prediction accuracy in high-confidence bins, with a 0.86 accuracy for predictions with confidences >0.9 (Fig. 2D). Similarly, applying the same binning strategy on 79 external Cas9 validation sequences showed that predictions with confidence >0.9 achieved a median accuracy of 0.95 (Fig. 2E).

We then applied CICERO-650M to the 54,539 Cas9 clusters for which we could not infer the PAM preference due to the lack of sufficient protospacers identified in phage and plasmid databases. To adhere to the same preprocessing routine as in training CICERO-650M, a length filter was applied, after which 50,308 sequences with a length between 200 and 1538 remained. These sequences were used as input for each trained fold of CICERO-650M, from which predictions and confidence scores were computed as the average across folds. This resulted in 32,016 predictions with confidence larger than 0.7, 17,453 predictions with confidence larger than 0.8, and 3,119 predictions with confidence larger than 0.9 (Fig. 3A). Using this dataset, we also constructed a phylogenetic tree of Cas9 protein clusters with high-confidence PAM predictions (confidence >0.7), providing a broad and systematic map of PAM diversity across 52,588 members of the Cas9 family (Fig. 3B).

**Figure 3.**
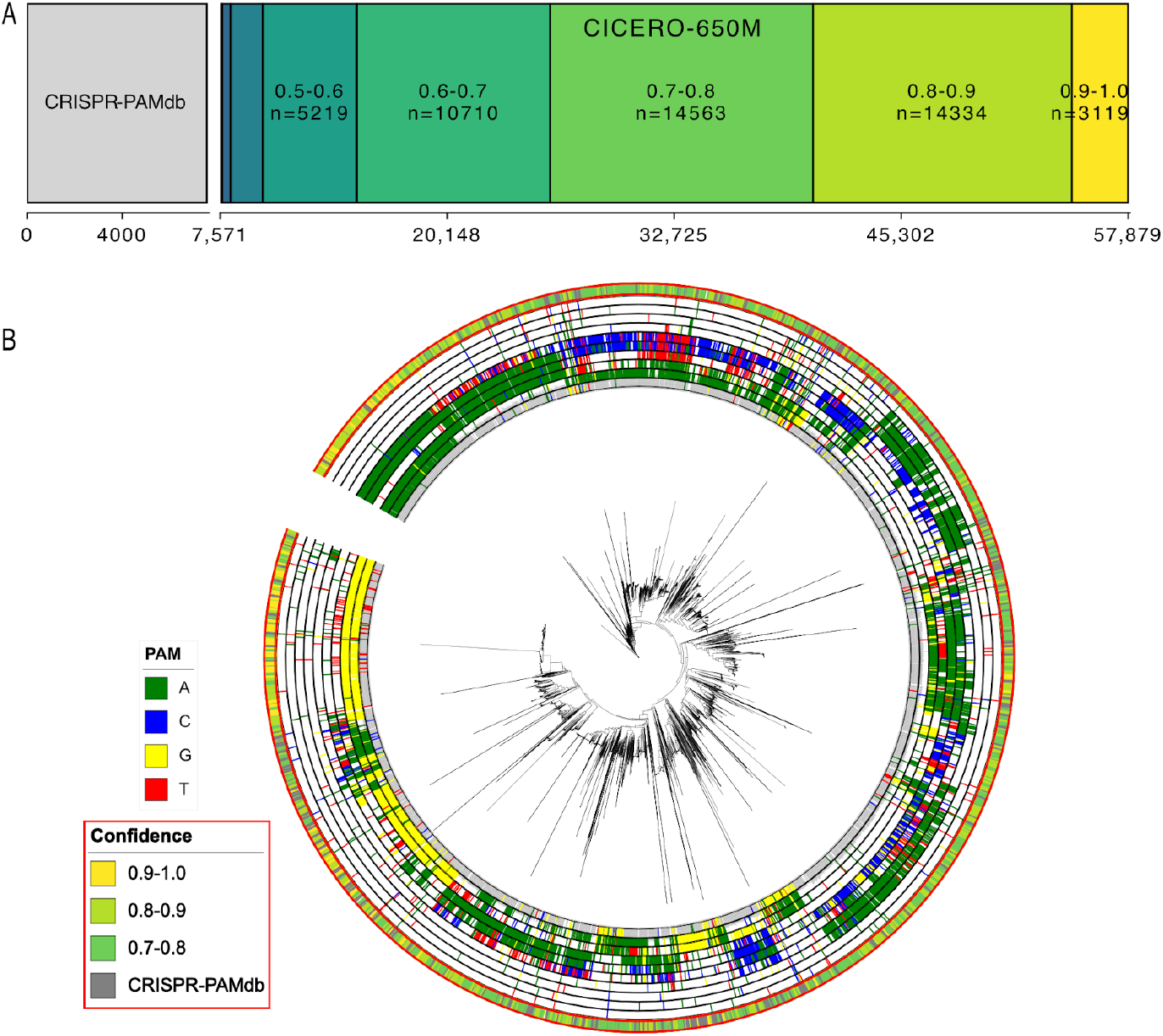
(A) Expansion of Cas9 PAM predictions using CICERO-650M. The initial 8,003 PAMs bioinformatically inferred from metagenomic datasets (gray) are extended to over 50,000 via CICERO-650M (colored). Each color represents a confidence bin of predicted PAMs. Notably, over 60% of the unlabeled Cas9 sequences received predictions with a confidence score above 0.7. (B) Phylogenetic tree of Cas9 protein clusters with consensus PAM inferred from metagenomic datasets or predicted with high-confidence (confidence score > 0.7) by CICERO. Annotations from the inner to outer rings represent the most likely nucleotide at each of the 10 PAM positions. The outermost ring indicates the source of the PAM prediction—inferred PAM (CRISPR-PAMdb) or CICERO—and, for CICERO-derived predictions, the associated confidence score.

## Discussion

In this study, we identified 62,542 unique Cas9 clusters from 3.8 million bacterial and archaeal genomes and screened 7.4 million phage and plasmid sequences to detect protospacer matches via spacer–protospacer alignment. This enabled the construction of CRISPR-PAMdb, a publicly available database containing 8,003 Cas9 clusters with inferred PAM preferences. To further expand PAM coverage, we developed CICERO, a machine learning model capable of directly predicting PAM profiles from Cas9 protein sequences. CICERO extended PAM annotations to an additional 50,308 Cas9 clusters, providing a systematic and scalable framework for exploring PAM diversity across the Cas9 protein family.

The accuracy of alignment-based PAM inference is inherently dependent on the diversity and abundance of unique spacers and protospacers available in current genomic datasets. A key strength of our study is the use of what is, to our knowledge, the most comprehensive collection of bacterial, archaeal, phage, and plasmid genomes assembled to date. This breadth is essential, as spacer and protospacer availability is shaped by ongoing host–invader co-evolution^25,26^. New spacers are regularly acquired in response to emerging threats, while older ones may be lost to preserve array compactness and efficiency. Similarly, protospacers in phage and plasmid genomes are frequently mutated or deleted to evade CRISPR-mediated interference.

To improve the robustness of inference, we clustered Cas9 proteins into homologous groups (>98% homology), allowing the aggregation of multiple unique spacers and protospacers for consensus PAM inference. While this approach increases statistical power, it may obscure differences in PAM preference between highly similar Cas9 variants. Nevertheless, when tested on a set of Cas9 variants with experimentally validated PAM preferences, we observed strong similarities to our inferred PAM profiles, suggesting that the CRISPR-PAMdb database will be a valuable resource for selecting Cas9 orthologs for genome editing experiments.

Compared to previous efforts^8,10^, our pipeline produced a higher number of Cas9 clusters with inferred PAM preferences, emphasizing the importance of leveraging comprehensive bacterial and archaeal genomic datasets, as well as comprehensive phage and plasmid sequence collections, for alignment-based PAM inference. Utilizing the taxonomic annotations provided by mOTUs4, we determined that 1,658,140 of 3,709,852 bacterial genomes (45%) and 23,352 of 62,669 archaeal genomes (37%) harbor Cas proteins. At the species level, 31,513 of 64,205 bacterial species (approximately 49%) were found to harbor Cas proteins. These findings are consistent with prior reports indicating that approximately 50% of bacterial genomes contain CRISPR systems. However, the lower-than-expected prevalence of Cas proteins in archaeal genomes—relative to historical estimates nearing 90%—may reflect gaps in phylogenetic coverage within our genome database or limitations of current Hidden Markov Models (HMMs) in detecting highly divergent or previously uncharacterized CRISPR–Cas variants in archaea. Alternatively, it may indicate the presence of distinct, non-CRISPR defense strategies in archaeal lineages^15,18,27,28^.

The potential of CICERO to predict PAMs from given Cas9 sequences massively extends the coverage beyond traditional alignment-based bioinformatics tools. By leveraging a protein language model (ESM2) and incorporating a confidence scoring mechanism, CICERO enables accurate and scalable prediction of PAM preferences directly from Cas9 sequence data. High-confidence predictions (e.g., confidence >0.9) corresponded to accuracies of up to 0.95 on benchmark datasets, offering a practical filter for prioritizing reliable predictions. Nonetheless, there is room for improvement. For example, future iterations of CICERO could benefit from expanded training datasets that include experimentally validated PAMs from more diverse Cas9 orthologs, particularly from underexplored phylogenetic clades. Moreover, CICERO is currently exclusively trained on Cas9 proteins, but the framework is inherently modular and training could be extended to other CRISPR effector families, such as Cas12 or Cas13.

Together, CRISPR-PAMdb and CICERO provide a powerful platform for the utilization of Cas orthologs targeting a diverse set of PAM sequences, paving the way for more flexible and targeted genome editing strategies.

## Methods

### Genomic Data Acquisition and CRISPR-Cas System Identification

3,747,151 isolated and metagenome-assembled bacterial and archaeal genomes were downloaded from the mOTUs4 database^12^. Among these, 37,298 were archaeal genomes. In addition, 25,371 archaeal genomes were retrieved from the European Nucleotide Archive (ENA) as of 2024-10-1^13^. To mine PAMs from mobile genetic elements, we established a reference dataset of phage sequences (N=7,978,168; Supplementary Table 2) from previously published bulk-metagenomic, viral-particle enrichment and viral isolate studies such as IMG-VR^20^ and added an in-house dataset of circular phages identified by mVIRs v.1.1.1 (standard parameters, minimal length of circular element is 2000 bp)^29^ as a circular element in the mOTUs-db assemblies^12^ and classified as “virus” by geNomad^30^ (v1.7.4, database version 1.7; *end-to-end* parameters: --disable-nn-classifcation, --sensitivity 7.5) (reference future downloadable file from mOTUs-db). The dataset was dereplicated using vClust v1.29^31^ (standard parameters for *prefilter* and *align* steps; --ani 1 --qcov 1 --tcov 1 --algorithm leiden for *cluster* step), which resulted in the total of 6,713,135 phage genomes. Additionally, we incorporated 699,973 plasmid genomes from IMG/PR^19^, resulting in a total of 7,413,108 mobile genetic elements.

The CRISPR array containing contigs were extracted from these genomes using MinCED (https://github.com/ctSkennerton/minced) and PILER-CR^32^. Results from both tools were merged, overlapping contigs were removed, and only contigs longer than 5,000 bp were retained. Protein-coding sequences within 20 kb upstream and downstream of CRISPR arrays in the contigs were predicted using Prodigal-gv v2.11.0^30^ and Cas variants were then identified by hmmsearch v3.4^33^ in these flanking regions using curated Cas Hidden Markov Models from CRISPRCasTyper^14^. We evaluated the pipeline using a manually curated CRISPR-Cas loci benchmark dataset comprising 1,106 genomes known to contain CRISPR-Cas9 systems^34^. Our pipeline successfully identified CRISPR-Cas9 systems in 1,012 of these genomes and recovered all curated Cas proteins—including, but not limited to, Cas9—in the majority of cases (Supplementary Fig. 1A).

### PAM Inference and Validation for CRISPR-Cas9 system

PAMs for the identified Cas9s were inferred by aligning associated spacers to protospacers in phage and plasmid genomes from our curated mobile genetic elements dataset (see above), discarding matches with more than four mismatches or gaps. Protospacers and their flanking regions (up to 10 nt on both sides) were aligned to derive consensus PAMs in the flanking region using PAMpredict^8^. To enhance PAMpredict’s accuracy, which depends on the number and diversity of input spacers, spacers from nearly identical Cas9 proteins were grouped using MMseqs2 (options: --min-seq-id 0.98 -c 1 --cov-mode 0)^35^ before analysis, improving PAM inference sensitivity. Spacer sequences belonging to nearly identical Cas9 proteins were also uniformly oriented before feeding to PAMpredict by aligning repeats, achieved through repeat clustering with cd-hit-est v4.8.1(options: -c 0.8, -s 0.75, -r 1)^36^. Redundant spacers were removed using cd-hit-est (options: -c 0.95, -s 1, -r 0), and Cas9 clusters with fewer than 10 spacers after dereplication were excluded from PAM inference. We benchmarked our pipeline using an external dataset of 79 Cas9 proteins with experimentally validated PAMs^21^ and achieved high inference accuracy for the 32 closely related proteins represented in CRISPR-PAMdb (Fig. 1E and Supplementary Fig. 1B).

### Machine Learning for PAM Prediction

In our experiments, we train CRISPR Cas9 PAM predictor model (CICERO) -an ML model that uses Cas9 protein sequences as input and predicts the associated PAM as an output. Our models use ESM2 as the backbone^11^ - a powerful protein language model that encodes information at the sequence level. Specifically, we employ the 650M parameter version in our approach CICERO-650M which encodes a Cas9 protein sequence and extracts a learned embedding representing a global contextual information across the entire protein, which is then fine-tuned during model training. In particular, we use the special [CLS] token, which is a meta token placed at the start of the input sequence for sequence representation. This is then passed to an output layer for PAM prediction - a fully-connected two-layer MLP with a rectified linear unit (ReLU) activation, a hidden dimension of 1280 and a dropout rate of 0.2. The MLP outputs a 10 × 4 matrix of raw PAM logits, corresponding to a fixed 10-length PAM prediction over ten nucleotide positions and four possible bases (A, T, G, C). See Fig. 2A for a visualization of the employed pipeline.

The training objective combines a standard cross-entropy loss on the predicted PAM logits, combined with an augmented cosine similarity loss at the information level, similar to Nayfach et al.^9^, to better align model predictions with biological PAM patterns. This augmented cosine similarity is also used as an evaluation metric for PAM predictions; it is considered a proxy for accuracy by quantifying how well the predicted PAM information content matches the target PAM profile inferred from metagenomic data by PAMpredict^37^. Intuitively, it captures both nucleotide agreement and the positional relevance of each base, encouraging the model to correctly identify high-information (i.e., biologically important) positions, while incorporating a fictitious “N” base to account for low-information regions where no specific base is strongly favored. All reported accuracy values throughout this paper refer to this metric.

We additionally implement and train a confidence network to enhance interpretability. In Phase 2 (Fig. 2A), we freeze both the finetuned protein language model backbone and the PAM-prediction head learned in Phase 1. We then attach a new confidence prediction head on top of the frozen [CLS] embedding. Only this confidence head is trained in this second phase to predict the empirical accuracy of the PAM head on each Cas9 input sequence. It is supervised with real-valued accuracy scores (computed per sequence) and optimized with an L2 loss. Intuitively, the confidence head learns to identify features in the finetuned, frozen [CLS] embeddings that predict errors of the PAM prediction head, allowing it to highlight inputs where the PAM prediction is likely unreliable.

For data processing, we used sequences for which a PAM was accurately inferred. As preprocessing, we apply length-based filtering by excluding sequences shorter than 200 amino acids and sequences exceeding the 99th quantile in terms of length (i.e., sequences longer than 1,538 amino acids). After filtering, we retain 7,571 pairs of sequences and their corresponding PAMs. For evaluation, we performed five-fold cross-validation using a stratified split setup with train/validation/test splits in each fold. Training is conducted using a learning rate of 1e-4 for both the protein language model backbone and the MLP head. The checkpoint with the best validation loss (evaluated on the validation split) is retained during training, which runs for 15 epochs, and is shown to be sufficient for our experiments. The confidence network is trained similarly on each fold for 15 epochs, with a learning rate of 3e-4. The models are first evaluated on the test split of each fold and subsequently on an external data set of 79 Cas9 sequences with experimentally determined PAMs^21^, which serves as additional ground truth.

To further investigate the performance characteristics of our models, we trained multiple CICERO variants with different ESM2 backbone sizes, ranging from 8M to 3B parameters. Evaluation was performed on the CRISPR-PAMdb test data, aggregated across all cross-validation folds. Supplementary Fig. 4A summarizes the accuracy of these models, both overall and above a confidence threshold of 0.8. Notably, we observe that performance increases consistently up to the 650M parameter model, but the 3B model shows diminished test accuracy, likely due to the limited size of the training dataset relative to model capacity. To evaluate calibration quality, we binned predictions by confidence quantiles and compared the predicted confidence to observed accuracy for each bin (Supplementary Fig. 4B), showing that CICERO-650M exhibits the best overall calibration, in particular for high-confidence regimes. Finally, we assessed accuracy stratified by input sequence length using equal-width bins over the observed length distribution (Supplementary Fig. 4C), which revealed that model performance is influenced by the input Cas9 protein length, especially for shorter sequences. This decrease in performance likely reflects limited data availability and increased variability in the shorter sequence regime of the CRISPR-PAMdb dataset^38^.

## Supporting information

Table S1. Predicted PAM results from both CRISPR-PAMdb and CICERO

Table S2. Data sources of phages included in our mobile genetic elements database

## Data and code availability

CRISPR-PAMdb was developed using Python for backend data processing and CICERO was implemented using PyTorch. The complete source code and accompanying documentation are available under an open-source license at https://github.com/Schwank-Lab/CRISPR-PAMdb and predicted PAM results from both CRISPR-PAMdb and CICERO can be downloaded from Supplementary Table 1.

## Acknowledgement

We thank the Schwank lab for their helpful discussions and feedback throughout the study. We also greatly appreciate the von Mering lab for their thoughtful input and for providing essential computational resources. This work was supported by core funding from ETH Zurich, the University Research Priority Program (URPP) Human Reproduction Reloaded of the University of Zurich, the Swiss National Science Foundation (SNSF) grant number 310030_185293, and the State Secretariat for Education, Research and Innovation–funded European Research Council Consolidator Grant (SERI-funded ERC-CoG) ‘GeneRepair’.

## Author information

These authors contributed equally: Tao Fang, Lea Bogensperger.

## Contributions

T.F.: Data curation, Formal analysis, Methodology, Project administration, Writing - original draft, Writing - review & editing;

L.B.: Formal analysis, Methodology, Project administration, Writing - original draft, Writing - review & editing;

L.F.: Software (snakemake pipeline), Code Review & Documentation, Writing - review & editing;

A.A.: Methodology, Supervision, Writing - review & editing; V.B.: Data curation, Resource, Writing - review & editing; Z.B.: Supervision, Writing - review & editing;

C.v.M: Resources, Supervision, Writing - review & editing; S.S.: Resources, Supervision,Writing - review & editing;

M.K: Conceptualization,Resources, Supervision, Funding acquisition, Writing - review & editing;

G.S.: Conceptualization, Resources, Supervision, Funding acquisition, Writing - review & editing.

## Supplemental Information

**Supplementary Figure 1.**
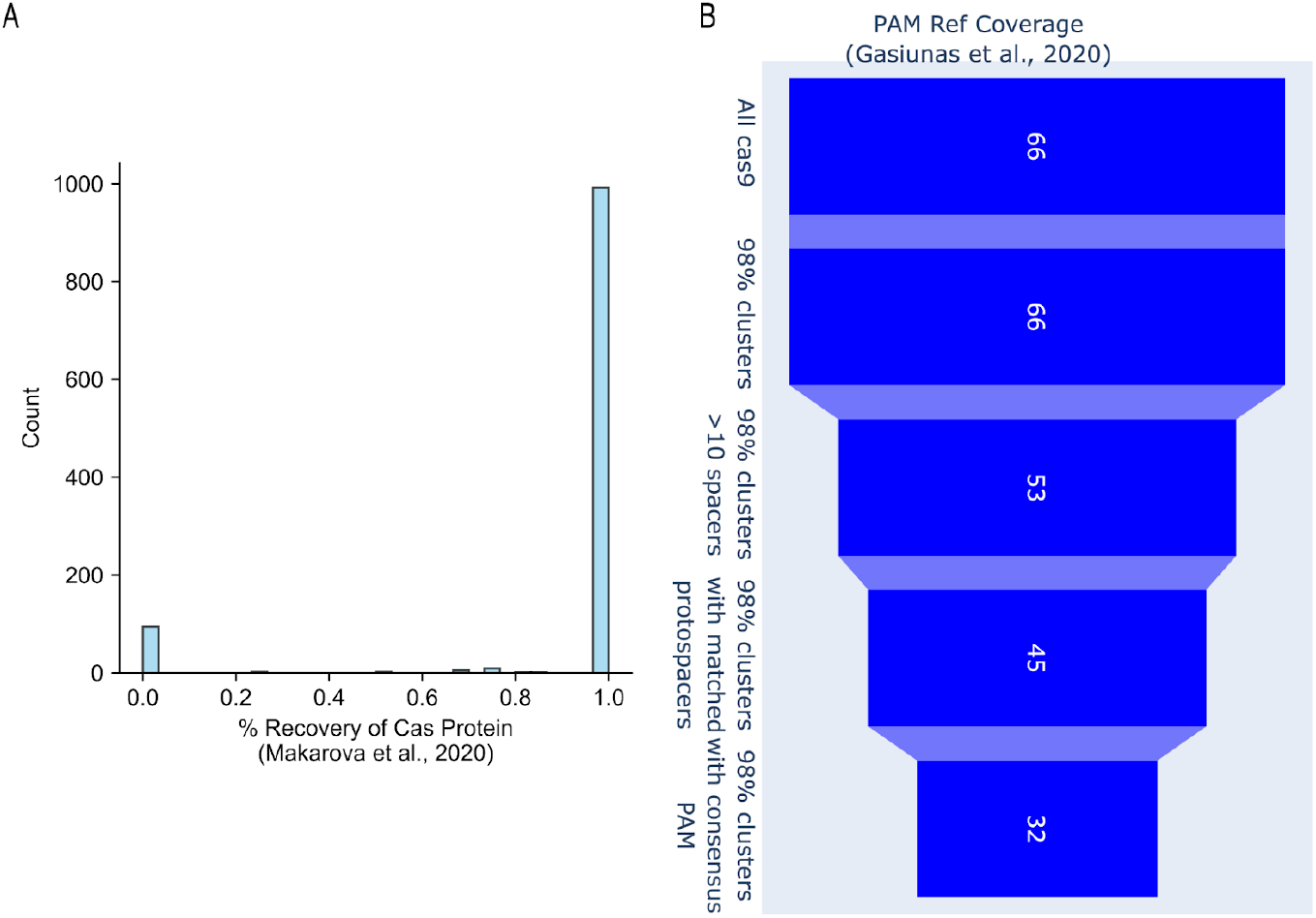
(A) Histogram showing the percentage of recovered Cas proteins (including, but not limited to, Cas9) from a manually curated CRISPR-Cas loci benchmark dataset consisting of 1,106 genomes known to contain CRISPR-Cas9 systems (See method section). (B) Coverage of experimentally validated reference PAMs (total of 79) at each pipeline stage, based on a 98% sequence similarity criterion.

**Supplementary Figure 2.**
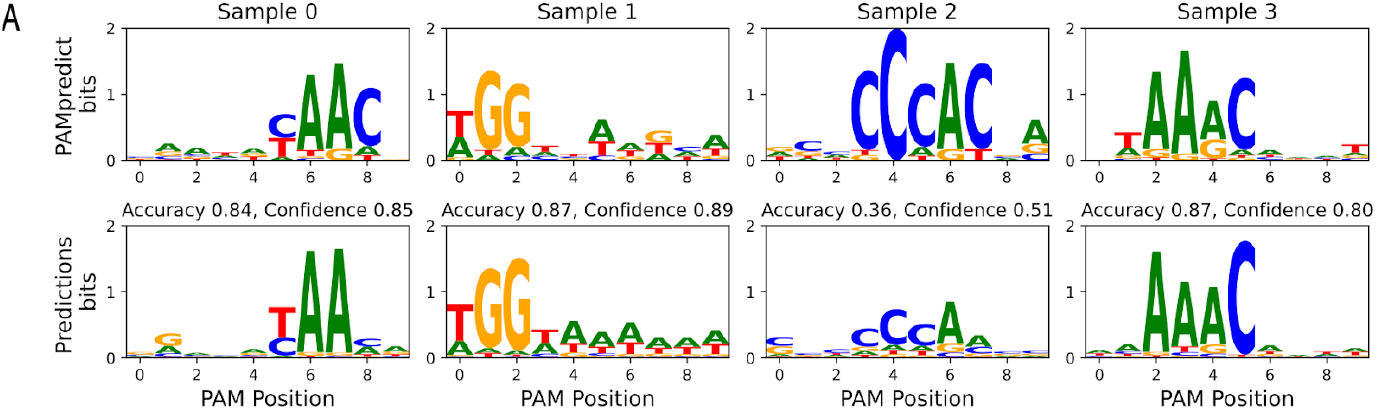
Examples for PAM logos obtained by alignment-based methods (top row) and predicted PAM logos (bottom row) for held-out test sequences from CRISPR-PAMdb using CICERO-650M.

**Supplementary Figure 3.**
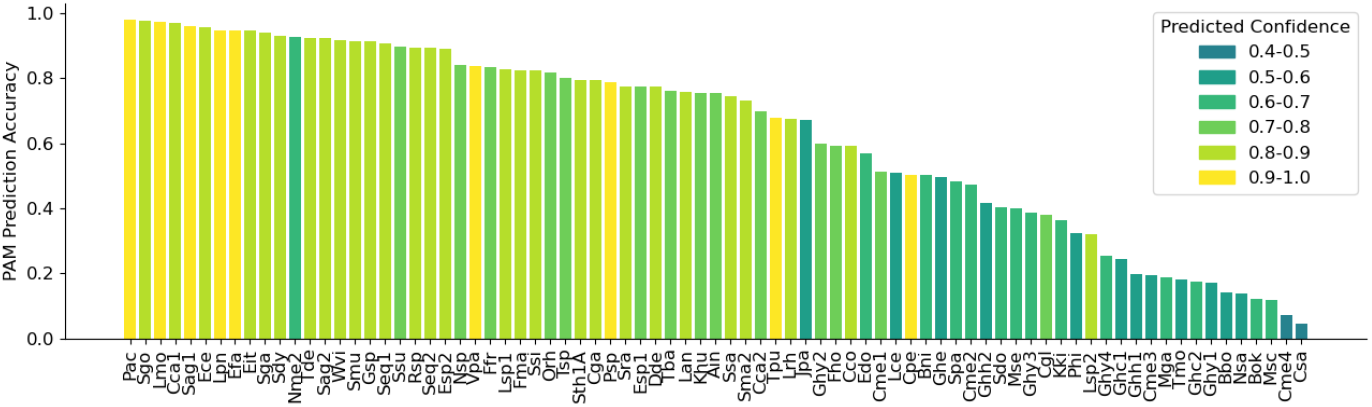
Per-sample predictions with a median accuracy of 0.75 for the 79 Cas9 sequences in the external validation data set. Note that each Cas9 sequence is colored according to the corresponding confidence obtained using the confidence network.

**Supplementary Figure 4.**
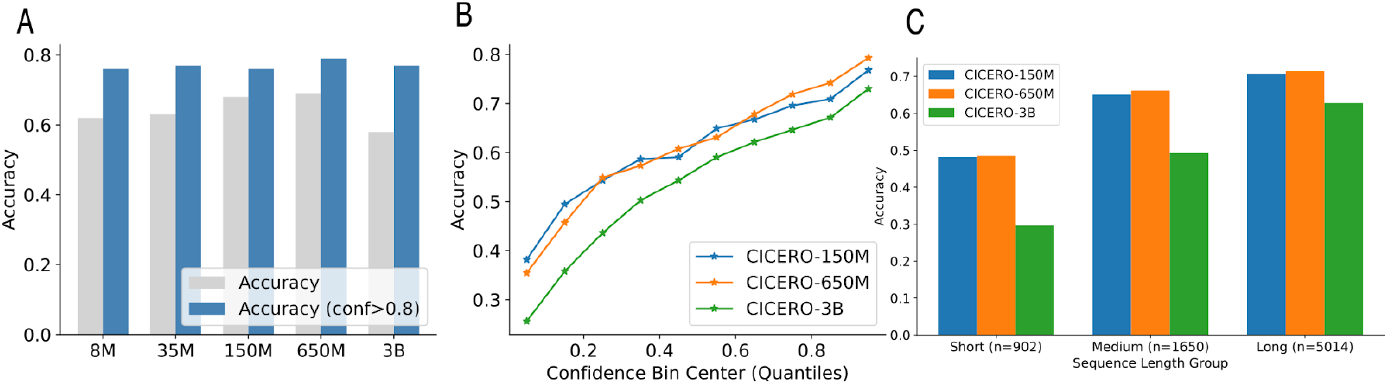
(A) Accuracy on the CRISPR-PAMdb test set aggregated across all folds. Gray bars show overall accuracy for each model size (CICERO-8M to CICERO-3B). Blue bars indicate accuracy when retaining only predictions with confidence scores above 0.8, demonstrating improved performance under high-confidence thresholds. (B) Calibration plots for the three largest CICERO models, showing actual accuracy within predicted confidence quantile bins. (C) Accuracies of the largest three models for stratified Cas9 sequence length. Short, medium and long sequence length refers to equidistant bins in the total sequence length range.

**Table S1.** Predicted PAM results from both CRISPR-PAMdb and CICERO

**Table S2.** Data sources of phages included in our mobile genetic elements database

